# Loss of PABPC1 is compensated by elevated PABPC4 and correlates with transcriptome changes

**DOI:** 10.1101/2021.02.07.430165

**Authors:** Jingwei Xie, Xiaoyu Wei, Yu Chen

## Abstract

Cytoplasmic poly(A) binding protein (PABP) is an essential translation factor that binds to the 3’ tail of mRNAs to promote translation and regulate mRNA stability. PABPC1 is the most abundant of several PABP isoforms that exist in mammals. Here, we used the CRISPR/Cas genome editing system to shift the isoform composition in HEK293 cells. Disruption of PABPC1 elevated PABPC4 levels. Transcriptome analysis revealed that the shift in the dominant PABP isoform was correlated with changes in key transcriptional regulators. This study provides insight into understanding the role of PABP isoforms in development and differentiation.

## 1. Introduction

Cytoplasmic poly(A) binding protein (PABP) is a key component of the translational machinery; it is critical for the closed loop formation of mRNA and stimulates mRNA translation into protein (Sonenberg and Hinnebusch 2009). PABP plays a direct role in 60S subunit joining and is integral to the formation of the translation initiation complex on the mRNA (Kahvejian, Svitkin et al. 2005). PABP also protects mRNA transcripts from decay (Coller, Gray et al. 1998).

Most structural and functional studies of cytoplasmic PABPs are based on PABPC1. PABPC1 is the most abundant of several cytoplasmic poly(A) binding proteins (PABPs) found in vertebrates and has been known for four decades (Blobel 1973). PABPC1 consists of four RNA-binding domains (RRM1-4) followed by a linker region and a conserved C-terminal MLLE domain. The RRM domains mediate the circularization of mRNA through the binding of the 3’ poly(A) tail and eIF4F complex on the mRNA 5’ cap (Imataka, Gradi et al. 1998, Deo, Bonanno et al. 1999, Kahvejian, Svitkin et al. 2005, Safaee, Kozlov et al. 2012). The linker region may promote the self-association of PABPC1 on mRNA although the molecular details of the interaction are unknown (Melo, Dhalia et al. 2003, Simon and Seraphin 2007). The C-terminus of PABPC1 contains a MLLE domain that mediates binding of a peptide motif, PAM2, found in many PABP-binding proteins (Xie, Kozlov et al. 2014).

Five other less abundant cytoplasmic PABPs exist in higher vertebrate. The PABP isoforms are believe to fulfill functionally distinct roles in vertebrate development (Gorgoni, Richardson et al. 2011). PABPC3 (tPABP or PABPC2 in mouse) is testis-specific (Kleene, Mulligan et al. 1998). PABPC4 (iPABP) is inducible in activated T cells (Yang, Duckett et al. 1995) and serves a critical role in erythroid differentiation (Kini, Kong et al. 2014). PABPC1L (ePABP) functions in oocytes and early embryos (Voeltz, Ongkasuwan et al. 2001, Seli, Lalioti et al. 2005, Guzeloglu-Kayisli, Pauli et al. 2008). PABPC1L is substituted by PABPC1 later in development, but remains expressed in ovaries and testes of adult (Vasudevan, Seli et al. 2006). PABPC4L and PABPC5 (Blanco, Sargent et al. 2001) lack the linker and MLLE domain.

There are additional binding sites for PABPC1 on mRNA transcripts in addition to the 3’ poly(A) tail. Gel shift assays show that the RRM domains of PABPC1 bind various RNA sequences other than poly(A); the RRM3-4 domains have broader specificity than RRM1-2 (Sladic, Lagnado et al. 2004). It was shown by CLIP-seq that only a low percentage (2.6%) of sequencing reads are pure poly(A) (Kini, Silverman et al. 2016). PABPC1 binds to an auto-regulatory sequence in the 5’-UTR of its own mRNA, and controls its own translation (de Melo Neto, Standart et al. 1995, Wu and Bag 1998, Hornstein, Harel et al. 1999). CLIP-seq study in mouse reveals that PABPC1 binds to a subset of A-rich sequences in 5’-UTR, besides predominant binding to 3’-UTR of mRNAs (Kini, Silverman et al. 2016). These PABPC1 interactions at 5’-UTR can impact and coordinate post-transcriptional controls on mRNAs (Kini, Silverman et al. 2016).

The functional specificity or redundancy of cytoplasmic PABPs is not well understood yet. There have been increasing interests in PABPC4 in recent years. PABPC4, a minor isoform of PABP, was first identified as an inducible protein in activated T-cells (Yang, Duckett et al. 1995). Depletion of PABPC4 interferes with embryonic development of *Xenopus laevis*, and cannot be rescued by isoforms PABPC1 or PABPC1L (Gorgoni, Richardson et al. 2011). PABPC4 plays an essential role in erythroid differentiation, and its depletion inhibits terminal erythroid maturation (Kini, Kong et al. 2014). Motif analyses of PABPC4 affected mRNAs reveal a high-value AU-rich motif in the 3’ untranslated regions (UTR) (Kini, Kong et al. 2014).

A major difficulty in studying the roles of PABP isoforms is the abundance of PABPC1 compared with the minor isoforms. To investigate specific or redundant roles of PABP isoforms, we disrupted PABPC1 in human cells with CRISPR/Cas9 gene editing system. An elevated level of PABPC4 compensated the loss of PABPC1, which suggested certain redundancy between the two isoforms. However, the transcriptome profile changed in PABPC4 elevated cells. Gene set enrichment analysis indicated that c-Myc was the most common gene in enriched pathways. Further, we showed correlated changes between PABP isoforms and c-Myc levels. These studies expand our understanding of cytoplasmic PABPs and suggest importance of a finely tuned network of PABP isoform usage.

## 2. Materials and methods

### 2.1 Cell culture and plasmids

Cells were cultured in DMEM supplemented with antibiotics and 10% fetal bovine serum. 10^5^ cells per well were plated in 24-well plate the day before transfection. 0.8 μg DNA plasmid was mixed with 2 μl Lipofectamine 2000 in Opti-MEM and then added to cells. After 24 h, cells were trypsin digested and split onto cover slides. pFRT/TO/FLAG/HA-DEST PABPC4 was from Thomas Tuschl (Addgene plasmid #19882) (Landthaler, Gaidatzis et al. 2008). DNA fragment expressing PABPC1 (NM_002568) or PABPC1Δ MLLE (1-542) were cloned into pCDNA3-EGFP between BamH I and Not I.

### 2.2 CRISPR/Cas9 genome editing

Target sequences were identified in PAPBC1 using CasFinder (http://arep.med.harvard.edu/CasFinder/) (Mali, Yang et al. 2013). hCas9 was from George Church (Addgene plasmid #41815). gBlocks expressing gRNA and the target sites were synthesized at Integrated DNA Technologies. The synthesized gBlocks were PCR amplified with primers gRNAforward/reverse for transfection into cells. 0.1×10^6^ HEK293T cells were transfected with 1 μg Cas9 plasmid, 1 μg gRNA, and 0.5 μg linealized NeoR gene fragment, using Lipofectamine 2000 as per the manufacturer’s protocols. Cells were split after 24 h, and selected in 400 μg/mL G418. Single cell-derived colonies were screened by western blotting for PABPC1 null mutations.

### 2.3 Primer and siRNA sequences

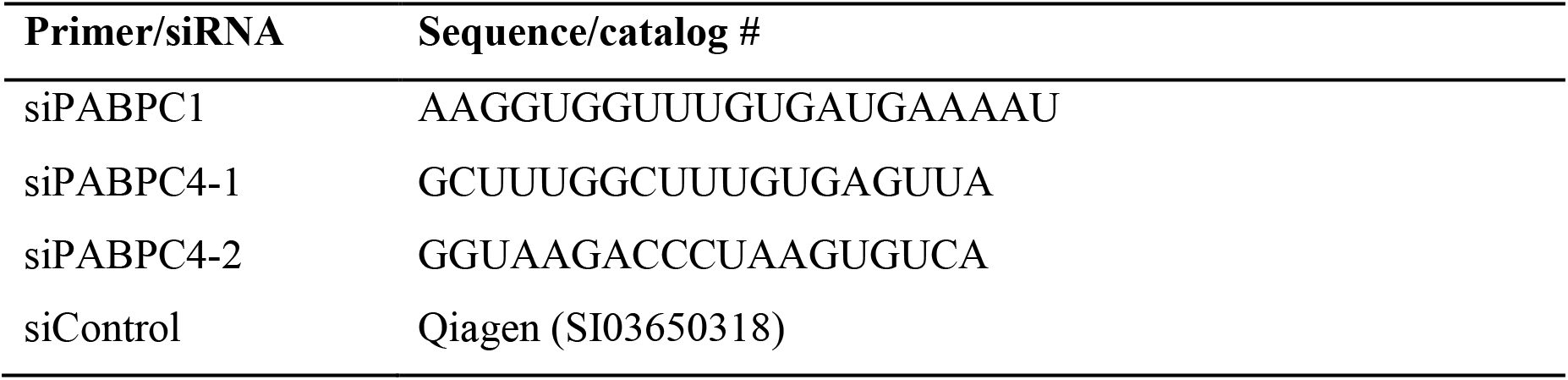
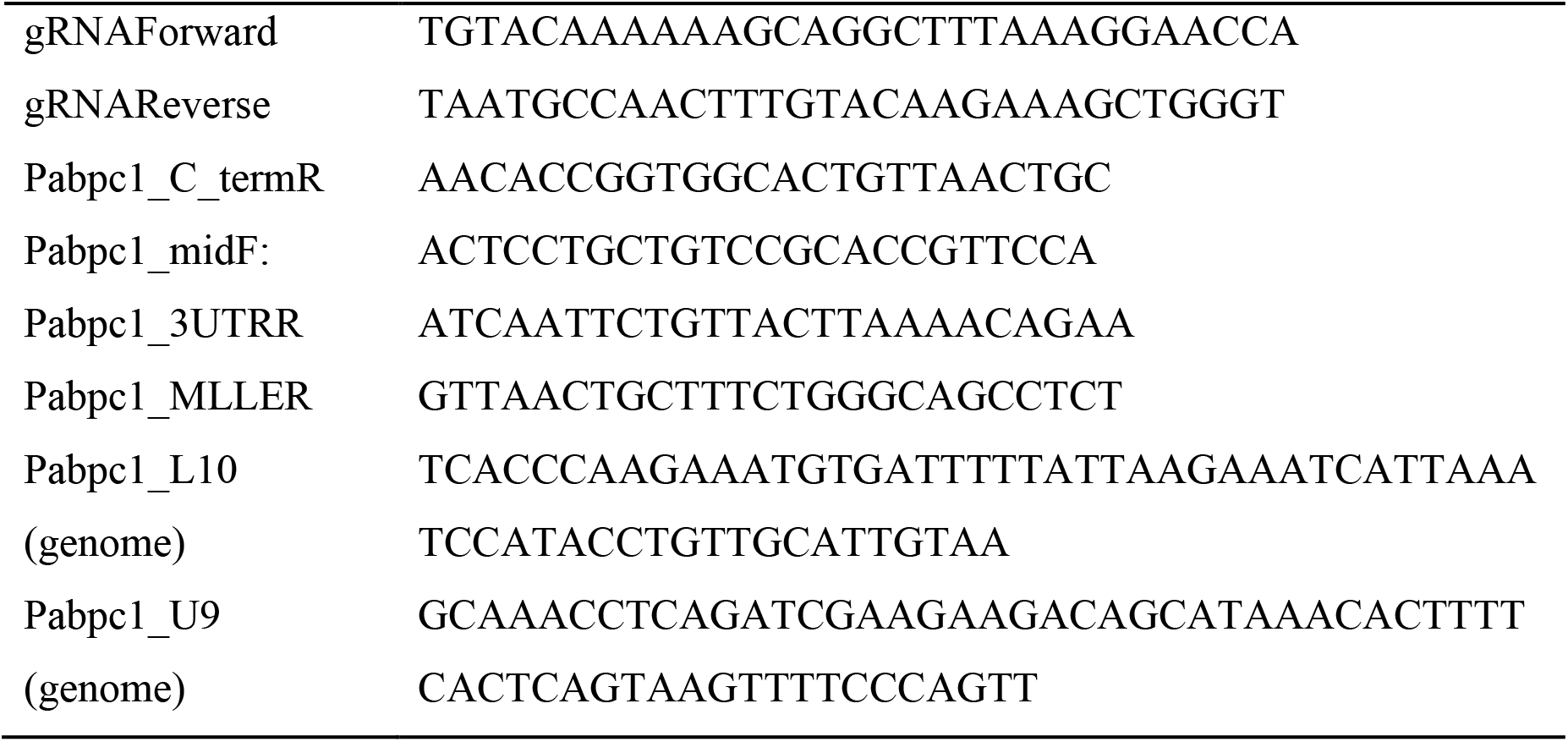

### 2.4 Genomic DNA extraction

Cells were pelleted and resuspended in 3 mL of TE buffer and 100 μL of 20% SDS. 20 μL of Proteinase K 20 mg/mL stock solution was added to mix gently by inversion and incubate overnight at 55 °C. 1 mL of saturated NaCl solution was added to the mixture. The solution was then precipitated overnight in 10 mL 100% EtOH (room temperature). DNA was transferred to 5 mL of 70% EtOH, and incubated overnight on rocker. DNA was then moved to a new Eppendorf tube and left air-dry. The DNA was dissolved in water and stored at −20 °C.

### 2.5 RNA extraction and cDNA library preparation

Total RNA was extracted from confluent 10-cm dishes using Trizol (Thermo-Fisher 15596-026) as per manufacturer’s instructions. RNA was air-dried and dissolved in distilled water. Dissolved RNA was further purified using Qiagen miRNeasy micro kit (Cat #217084). RNA from replicates of HEK293 or clone-c1c4 cells were aliquoted and stored at −80 °C for later quantitative RT-PCR analysis or sent for mRNAseq library preparation at Genome Quebec Innovation Center.

### 2.6 RNA-seq and Differential gene expression analysis

Pair-ended RNA sequencing with read lengths of 100 bases was performed at the Genome Quebec Innovation Center using the Illumina Hiseq 2000 sequencer. Reads were trimmed from the 3’ end to have a phred score of at least 30. Illumina sequencing adapters were removed from the reads, and all reads were required to have a length of at least 32. Trimming and clipping were done with Trimmomatic (http://www.usadellab.org/cms/index.php?page=trimmomatic). The filtered reads were aligned to reference genome b37. The alignment was done with the combination of tophat/bowtie software to generate a Binary Alignment Map file (Trapnell, Pachter et al. 2009). Read counts were obtained using HTSeq as input. The differential gene expression analysis was done using DESeq (Anders and Huber 2010) and edgeR (Robinson, McCarthy et al. 2010) Bioconductor package. The results of the differential transcript expression analysis were generated using cuffdiff. FPKM values calculated by cufflinks were used as input (Trapnell, Roberts et al. 2012).

### 2.7 Preranked gene set enrichment analysis

Differential genes from Deseq were ranked by their fold of changes (FC) and p-values (Ranking score = log_2_FC ×(−log_10_(p-value)) (Supplemental Table 3) (Plaisier, Taschereau et al. 2010). The ranked gene list was used as input for GSEA and leading edge analysis (Mootha, Lindgren et al. 2003, Subramanian, Tamayo et al. 2005). The detailed GSEA parameters were as follows: the number of permutations is 1000, and the permutation type was configured to the gene set.

### 2.8 Immunofluorescence and confocal microscopy

Cells were fixed with 4% PFA in PBS, and penetrated by cold methanol (−20 °C) or 0.1% Triton X-100 in PBS for 10 min. Cells were blocked with 5% goat serum (Millipore S26) in PBS for 1 h. Then cells were incubated in PBS, supplemented with anti-PABPC1 (Santa Cruz sc32318, 1:200) and anti-PABPC4 (PTGlab AP-14960, 1:200). Cells were washed in PBS three times, before incubation with second antibodies conjugated with Alexa488 or Alexa647 (Sigma-Aldrich A31620, A31628, A31571) at 1:200 – 1:500 dilutions. DAPI (Roche) was added to the first wash at 0.5 μg/ml for 10 min. Cover slides were mounted in ProLong Gold anti-fade reagent (Life Technology P36930). Images were collected on a Zeiss LSM 310 confocal microscope in the McGill University Life Sciences Complex Advanced BioImaging Facility (ABIF).

### 2.9 Quantitative RT-PCR

Total RNA was extracted from cells with Trizol (Life Technology). cDNA libraries were prepared using SuperScript First-Strand Synthesis System for RT-PCR (Life Technology). Validated Taqman assays were purchased for quantification of GAPDH (Applied Biosystems Hs 02758991), c-Myc (Applied Biosystems Hs00153408), and 18sRNA (Applied Biosystems Hs 99999901). qRT-PCR were run and analyzed in Stepone Plus PCR system (Applied Biosystems).

### 2.10 Western blotting

Protein samples were heated at 95°C and separated by SDS-PAGE. Proteins were then transferred to PVDF membrane (Millipore) in Tris/Glycine buffer with 20% methanol in cold room. PVDF membrane was blocked in TBST (pH 7.5), containing 0.05% Tween-20 and 5% skim milk powder or bovine serum albumin. The membrane was then incubated with primary antibodies, including anti-PABPC1 (Abcam ab21060, Cell signaling 4992 or Santa Cruz sc32318 1:1000), anti-PABPC4 (Abcam ab76763), anti-c-Myc (Santa Cruz sc 40), anti-tubulin (Sigma-Aldrich T9028 1:5000) and anti-GFP (Clontech 632381 1:2000). The membrane was then washed three times in TBST and incubated with goat-anti-rabbit (Jackson ImmunoResearch 111-035-046 1:5000) or goat-anti-mouse (Jackson ImmunoResearch 115-035-071 1:5000) for 0.5 h, washed again, developed with Amersham ECL prime kit (GE healthcare RPN2236), and imaged on an Alpha Innotech imaging system.

## 3. Results

### 3.1 PABPC4 compensates partial loss of PABPC1 in HEK293

PABPC1 is the predominant isoform of cytoplasmic PABP in cells. We chose to edit endogenous PABPC1 to check for changes in expression of other isoforms. We selected two target sites in exon 10 of the *Pabpc1* locus, and incorporated the sequences into gBlocks expressing guide RNA scaffolds (Fig. S1). The two scaffold RNAs were separately transfected into HEK293, together with Cas9 plasmid and a linearized NeoR gene for antibiotic selection. Random deletion, insertion or mutation was introduced around the targeted sites. Single cell colonies were screened with an antibody recognizing the C-terminal of PABPC1, for mutant cell-lines expressing PABPC1 mutants altered after the target sites. About 20% of the colonies had insertion or deletions leading to a shortened or disrupted PABPC1 (Fig. 1A). Two typical cell-lines, clone-c1 expressing an exon-skipped PABPC1 (Fig. 2 and Fig. S2), and clone-c1c4 (from target sequence 2) expressing a truncated PABPC1 (Fig. 1A&D) were selected for genomic DNA sequencing (Fig. 1C). Deletion of 3 base pairs in clone-c1 resulted in skipping of the whole exon 10 (Fig. S2). The 2 base pair insertion in clone-c1c4, led to truncated *Pabpc1* mRNA (Fig. 1D) and a corresponding PABPC1 protein truncates due to early termination after the two base pair insertion (Fig. 1B and Fig. 2). Clone-c1c4 was selected for further study, as its PABPC1 protein was totally disrupted and greatly decreased. An approximate two-fold elevation of PABPC4 in protein and mRNA levels was observed in clone-c1c4, where the PABPC1 is truncated and decreased (Fig. 1A & E). The major PABP isoform in clone-c1c4 is PABPC4, instead of PABPC1 as in HEK293.

**Figure 1.**
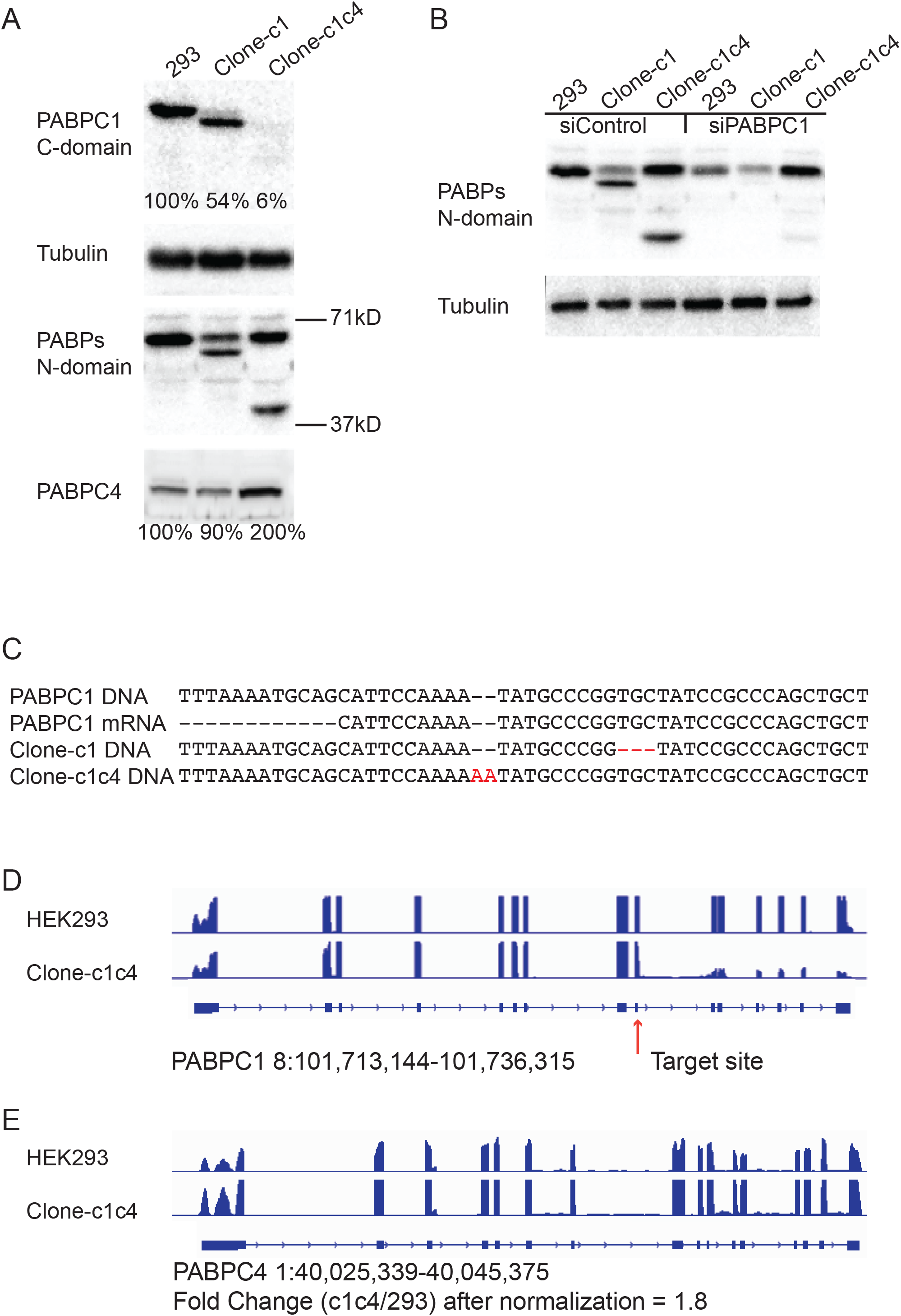
PABPC1 disruption in HEK293 cells by CRISPR/Cas9 based genome editing is compensated by elevated PABPC4. (A) CRISPR/Cas9 genome editing of PABPC1 generated multiple mutations. Antibody recognizing C-terminal MLLE domain of PABPC1 (PABPC1 C-domain, Santa Cruz 32318) labels null mutations of PABPC1. PABPs N-domain antibody (New England Biolabs 4992), which recognizes multiple cytoplasmic PABP, reveals PABP isoform distribution. Tubulin is stained as a loading control. Anti-PABPC4 shows an elevated level of PABPC4 protein in clone-c1c4, when PABPC1 is truncated to about 40 kD and less expressed. Percentages are relative protein levels after normalized to tubulin. (B) PABPC1 was knocked-down by siRNA, in 293, clone-c1, or clone-c1c4 cells. The decreased lower bands in clone-c1 and clone-c1c4 were PABPC1 mutations. Sequences of siRNAs used are listed in materials section. (C) Genomic DNA sequences at *Pabpc1* gene loci of 293, clone-c1, and clone-c1c4. Deletion of three base pairs in clone-c1 leads to skipping of exon 10 in mRNA (Fig. S2) and a shorter PABPC1 protein (Fig. 2). Clone-c1c4 has a two base pair insertion, which results in significant reduction of mRNA reads after the targeted region (D). Genomic DNA was extracted and amplified with gene specific primers. PCR products were gel purified and cloned into PCR2.1 vector for sequencing. Only one sequence was read in multiple clones, indicating the homogeneity of cell-lines. (D & E) Wiggle tracks showing representative read alignment of *Pabpc1* or *Pabpc4* genes in HEK293 or clone-c1c4 cells. Track files were generated from the aligned reads using BedGraphToBigWig.

**Figure 2.**
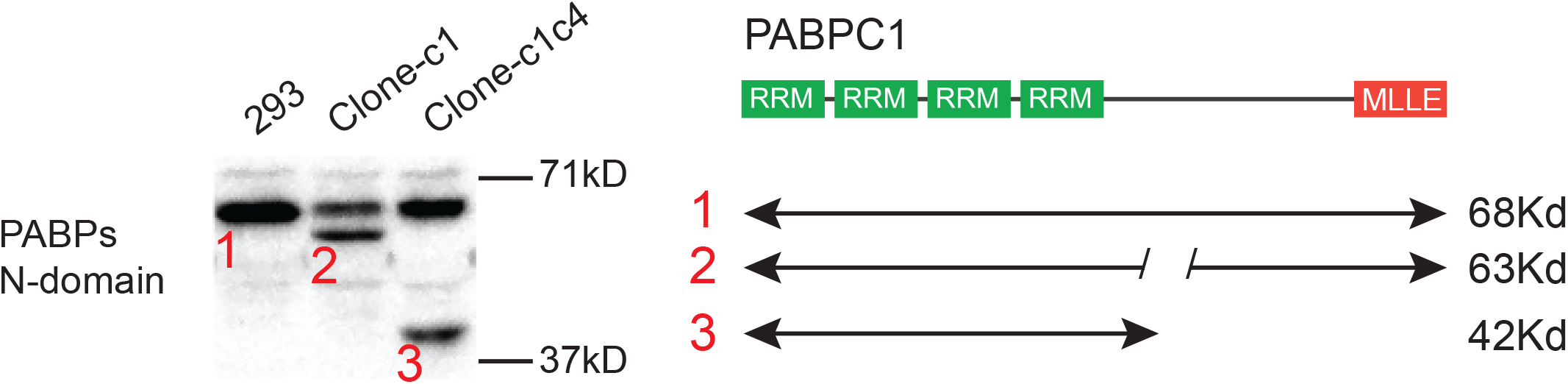
Description of PABPC1 protein in HEK293, clone-c1, and clone-c1c4. Wild-type PABPC1 is 68 kD in molecular weight. A shortened form of PABPC1 was generated in clone-c1 with residues 447-483 deleted. The mRNA sequence of clone-c1 is shown in Figure S2. In clone-c1c4, the major PABPC1 was about 42 kD in agreement with the size of the major mRNA species (Fig. 1E).

### 3.2 Overexpression of PABPC1 represses elevated PABPC4 in clone-c1c4 cells

The shift of dominative PABP isoform from PABPC1 to PABPC4 did not change cell proliferation (data not shown) or morphology (Fig. 6B). This suggests that the two isoforms are redundant in maintaining basic cellular activities. We then asked whether the elevated PABPC4 in clone-c1c4 was reversible by expression of PABPC1. Clone-c1c4 cells were overexpressed with PABPC1 or PABPC1ΔMLLE (Fig. 3A). PABPC1ΔMLLE overexpression repressed PABPC4 more in clone-c1c4 cells. It is not clear why deletion of the MLLE domain enhances the repression activity of PABPC1. However, it indicates that the repression comes from the RNA binding ability of the RRM domains.

**Figure 3.**
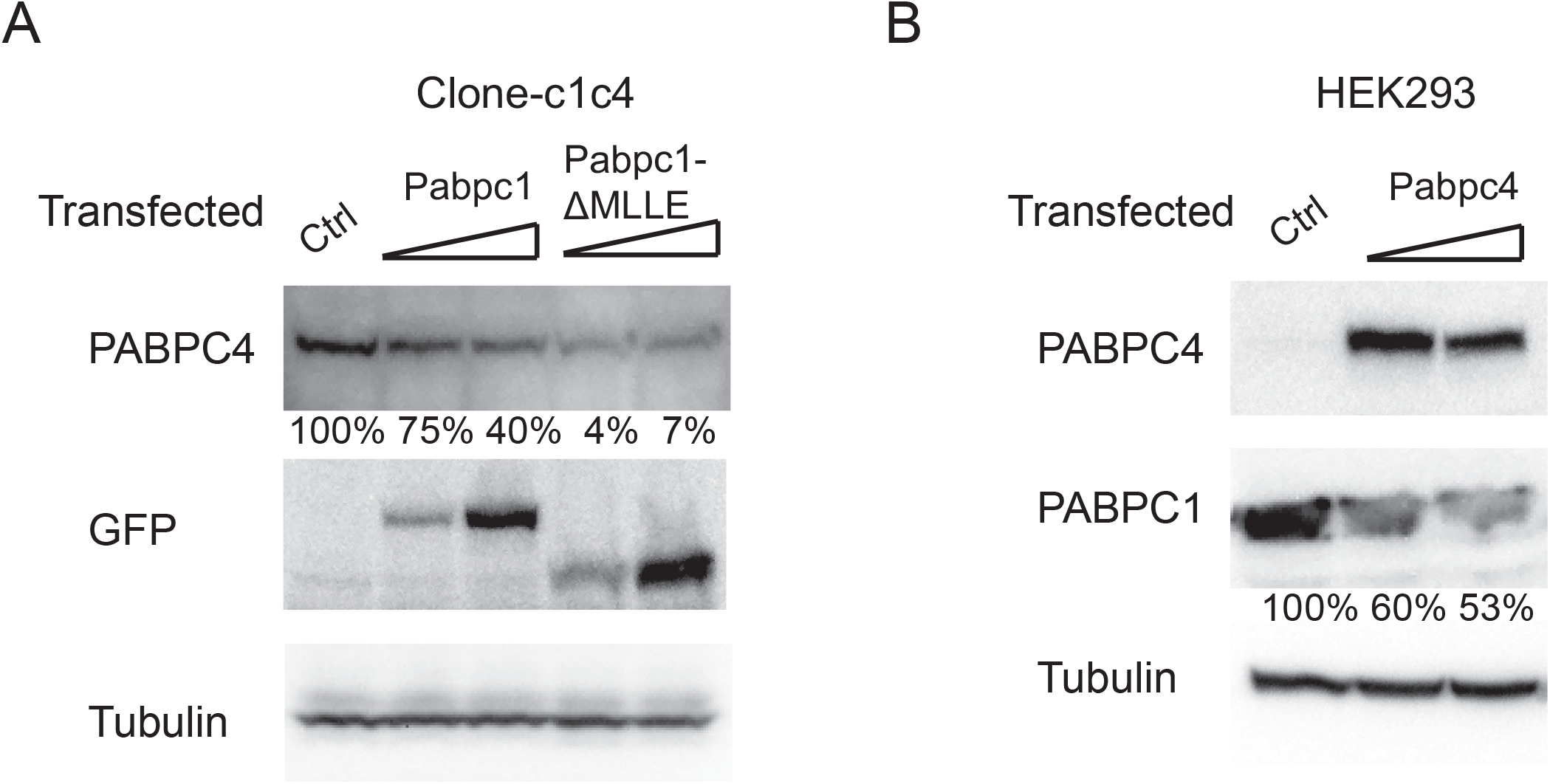
Mutual repression of PABPC1 and PABPC4. (A) Over-expression of pCDNA3-EGFP (Ctrl), pCDNA3-PABPC1-EGFP, or pCDNA3-PABPC1ΔMLLE-EGFP in clone-c1c4 cells. Western blotting shows decrease of PABPC4 protein level in clone-c1c4 cells, due to over expression of PABPC1-EGFP or PABPC1ΔMLLE-EGFP. (B) Over-expression of PABPC4 (construct in materials) in HEK293 reduces endogenous PABPC1 protein.

### 3.3 Overexpression of PABPC4 represses endogenous PABPC1 in HEK 293

We next overexpressed PABPC4 in HEK293, and found a reduction of endogenous PABPC1 (Fig. 3B). This further confirms the redundancy of isoforms PABPC1 and PABPC4. The total amount of PABP isoforms may be well regulated for cellular operations. Thus the elevation of PABPC4 in clone-c1c4 cells is due to loss of PABPC1.

### 3.4 Differential gene expression analysis in clone-c1c4 cells

The clone-c1c4 cell-line offered us a platform to study PABP isoform specific functions. To investigate the effects of PABP isoform usage shift on the transcriptome, we submitted clone-c1c4 and HEK293 cells for RNA-seq. Read counts were obtained using HTSeq. Differential gene expression analysis was done with edgeR (Robinson, McCarthy et al. 2010) and DESeq (Anders and Huber 2010) R bioconductor packages. Differential expressed (DE) genes were ranked according to the p-values. The gene expression heat map (Fig. S3) indicated profile changes in clone-c1c4 cells. The top 300 DE genes were labeled red on MA plot (Fig. 4A). Most of the top 300 DE genes were relatively highly expressed according to the counts. The top ten DE genes were shown in table (Fig. 4B). Representative wiggle track views of the ten genes displayed mRNA profiles, confirming the DE gene calling (Fig. 4C).

**Figure 4.**
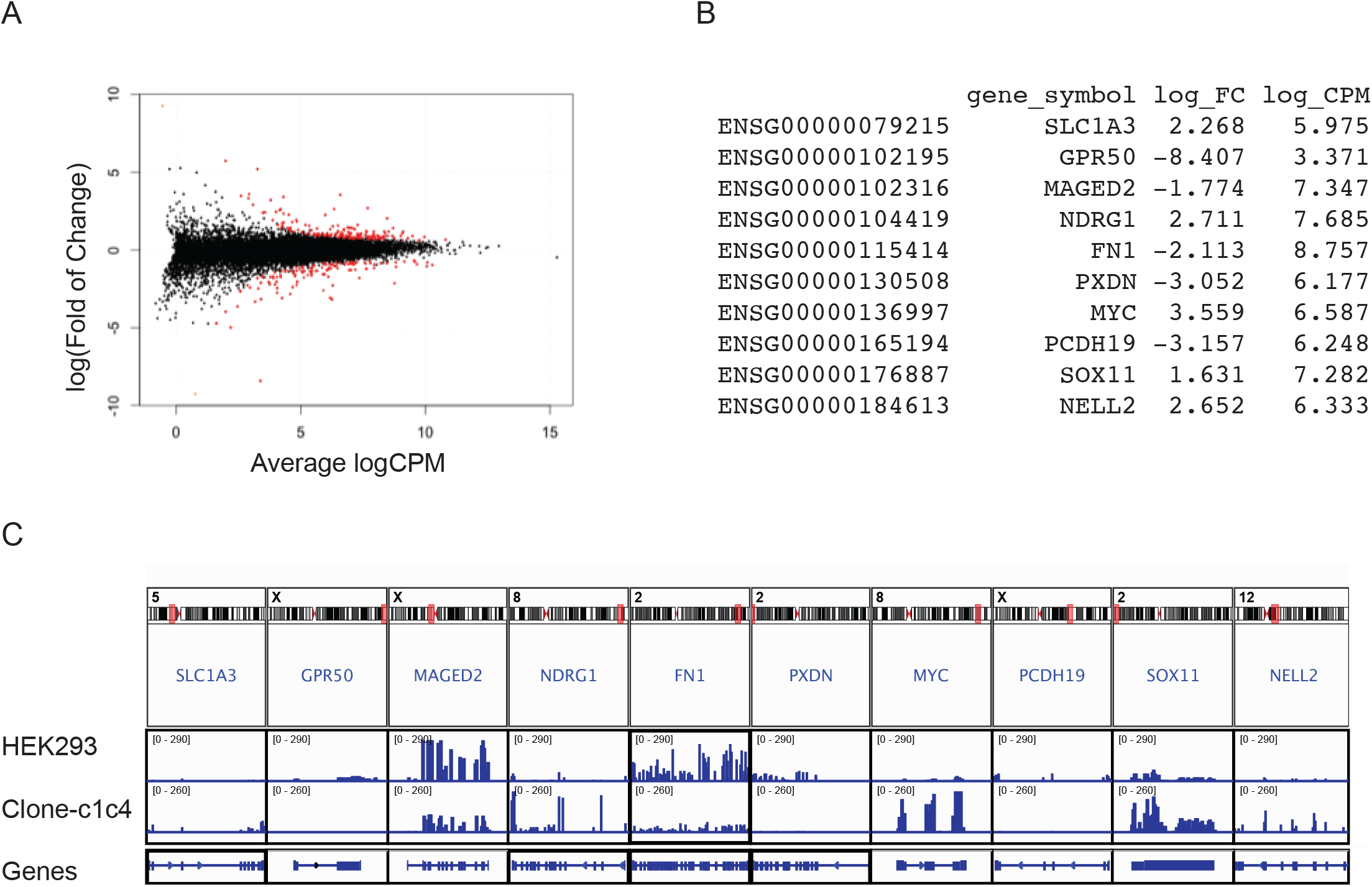
Differential gene expression in HEK293 and the modified clone-c1c4 cells. (A) MA plot of differential expression magnitude (log2 of fold change (clone-c1c4/HEK293) versus expression levels (log2 of counts per million). The red dots are the top 300 differential genes ranked by p-values. (B) The top 10 differential genes. Ensembl gene id, gene symbol, log2 (fold of changes), and log2 (counts per million) are shown in the table. (C) Representative view of the mRNA profile of the top 10 differential genes. The figure is generated with Integrative Genomics Viewer.

### 3.5 c-Myc is central to differential gene expression in clone-c1c4

To infer biologically important genes underlying the transcriptome changes, we used Gene Set Enrichment Analysis to examine the enrichment of 50 hallmark signature gene sets (Liberzon, Birger et al. 2015). Enriched gene sets were plotted against their normalized enrichment scores. Gene sets with p-values lower than 0.10 are marked red (Fig. 5A). The leading edge subset analysis of the most enriched gene sets revealed c-Myc as the most overlapped gene (Fig. 5B and Fig. S4). The increase of c-Myc in clone-c1c4 was confirmed by western blotting and qRT-PCR (Fig. 5 C & D).

**Figure 5.**
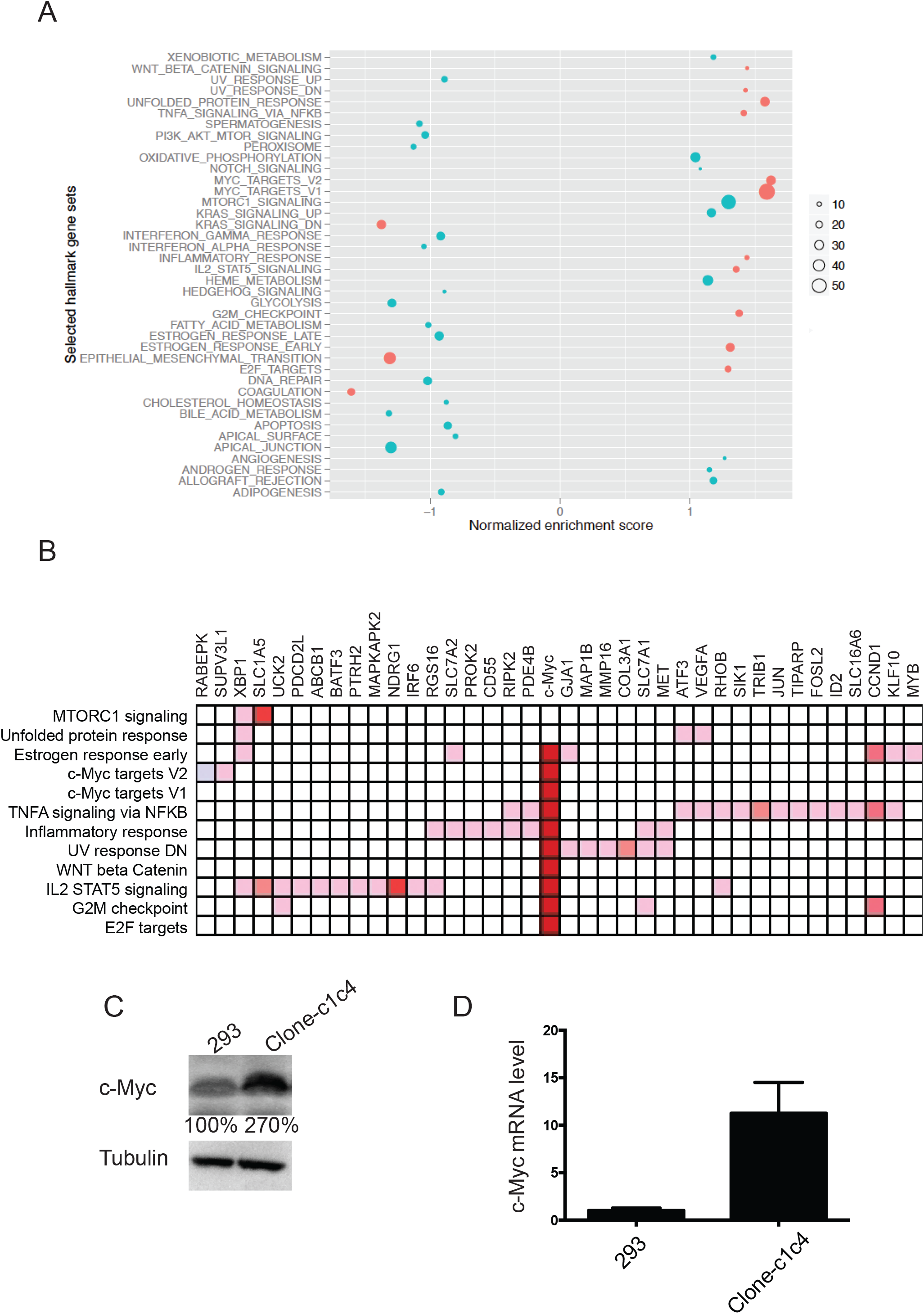
Gene expression in clone-c1c4 cells. (A) Gene set enrichment analysis of selected hallmark gene sets (Subramanian, Tamayo et al. 2005). The size of circles indicates the number of significant genes. Gene sets with p-values lower than 0.1 are labeled red. (B) Leading edge overlap for the modified clone-c1c4 cells. The significantly enriched gene sets from (A) are aligned to indicate common genes. c-Myc is the most significant overlapped gene. Other overlapped genes are shown in figure S4. The intensity of color indicates fold of changes in expression. Only a subset of the genes are displayed for visibility. (C) Increased c-Myc protein in clone-c1c4 cells. The percentage indicates relative quantifications of c-Myc protein after normalization. Cells were lysed in SDS-loading buffer and boiled. c-Myc protein level was probed by antibody (Sant Cruz sc-40). Tubulin is used as loading control. (D) Increased c-Myc mRNA level. Taqman assay (Applied Biosystems Hs 00153408) reveals an increase of about 10-fold in c-Myc mRNA in clone-c1c4. GAPDH (Applied Biosystems Hs 02758991) is used as loading control.

**Figure 6.**
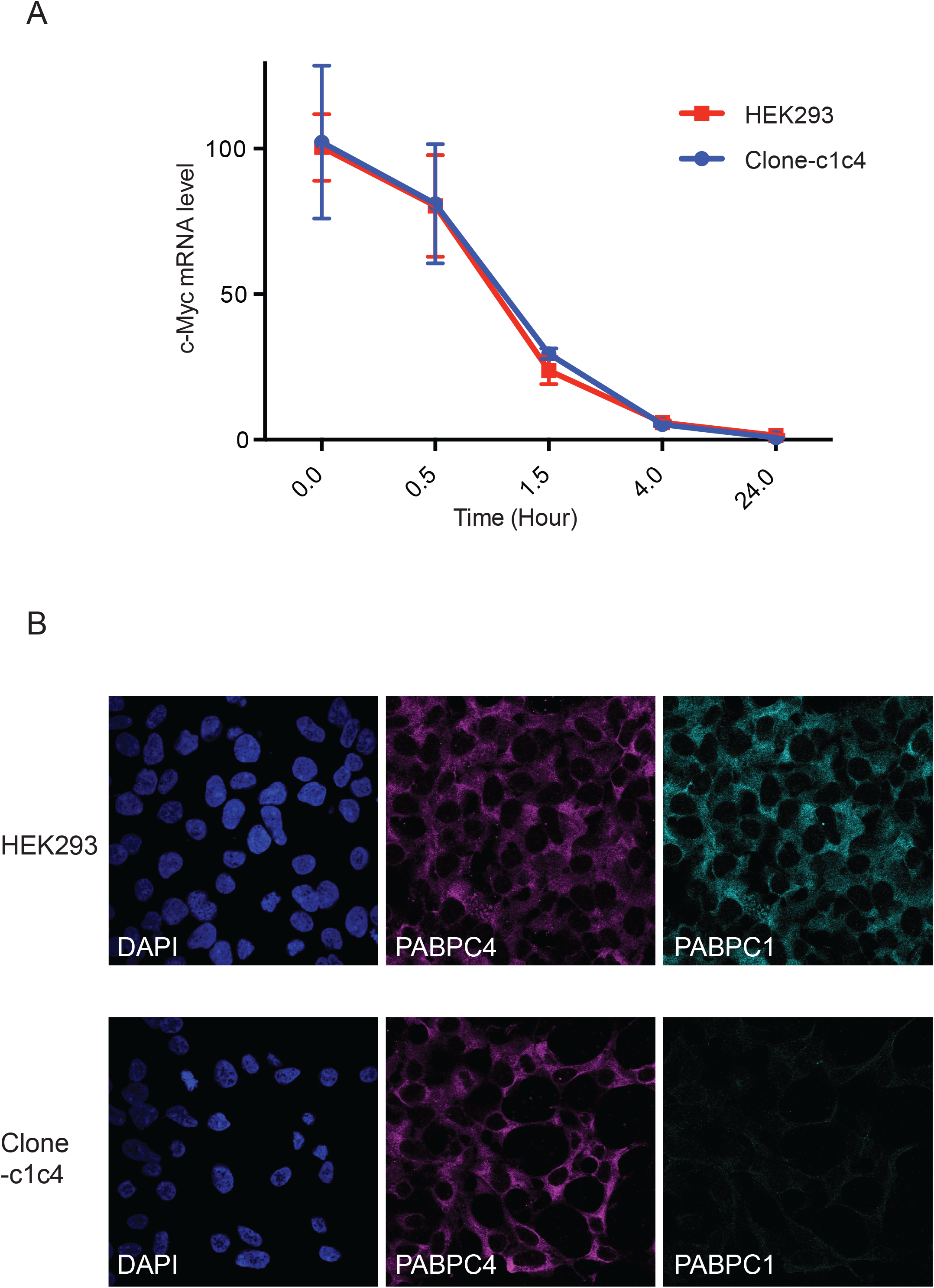
(A) c-Myc mRNA half life is similar in HEK293 and clone-c1c4. Cells were treated by (10 μg/ml) antinomycin-D for indicated time before Trizol extraction of total RNA. Taqman assays were used to measure c-Myc levels normalized to 18s RNA (Applied Biosystems Hs 99999901). The c-Myc mRNA levels of cells without treatment are normalized to 100 for comparison. (B) PABPC4 in clone-c1c4 is predominantly cytoplasmic. HEK293 or clone-c1c4 cells were stained with DPAI, anti-PABPC4 (Abcam ab76763), and anti-PABPC1 (Abcam ab21060). The PABPC1 antibody recognizes the C-terminal tail of PABPC1.

### 3.6 c-Myc mRNA half life is not changed in clone-c1c4

To check whether the increased c-Myc mRNA level in clone-c1c4 was transcriptional or post-transcriptional, we treated HEK293 or clone-c1c4 cells with 10 μg/mL actinomycin-D to inhibit new transcription. Samples were collected at different time points to determine c-Myc mRNA half-life by quantitative RT-PCR. C-Myc mRNA levels measured by taqman assay were normalized to 18s RNA (Fig. 6A). This suggests that the increase of c-Myc mRNA level is transcriptional.

### 3.7 PABPC4 is predominantly cytoplasmic in clone-c1c4

Although role of PABPC4 in transcriptional control is not known, there is increased nuclear distribution of PABPC4 in response to stress (Burgess, Richardson et al. 2011) or PABPN1 depletion (Bhattacharjee and Bag 2012). We stained PABPC4 in HEK293 and clone-c1c4 cells, and found PABPC4 are predominantly cytoplasmic in both cell-lines (Fig. 6B). Thus the increased PABPC4 in clone-c1c4 doesn’t lead to significant nuclear relocation.

### 3.8 Correlation of PABPC4 and c-Myc changes

We then knocked-down *Pabpc4* expression in clone-c1c4 cells with siRNAs. c-Myc mRNA and protein levels decreased correlated to PABPC4 depletion (Fig. 7A & B). We asked whether substitution of PABPC1 by PABPC4 led to the c-Myc increase. Overexpression of PABPC1-GFP in clone-c1c4 repressed endogenous PABPC4 protein (Fig. 3A). However, the increased PABPC1-GFP in clone-c1c4 did not decrease the c-Myc mRNA level. We reasoned that it might take longer for the PABPC1 to indirectly affect the c-Myc mRNA level, or the relatively large GFP tag might interfere with certain PABPC1 functions. Nonetheless, the relative usage of PABPC4 isoform correlates with the c-Myc level and can potentially affect the transcriptome.

**Figure 7.**
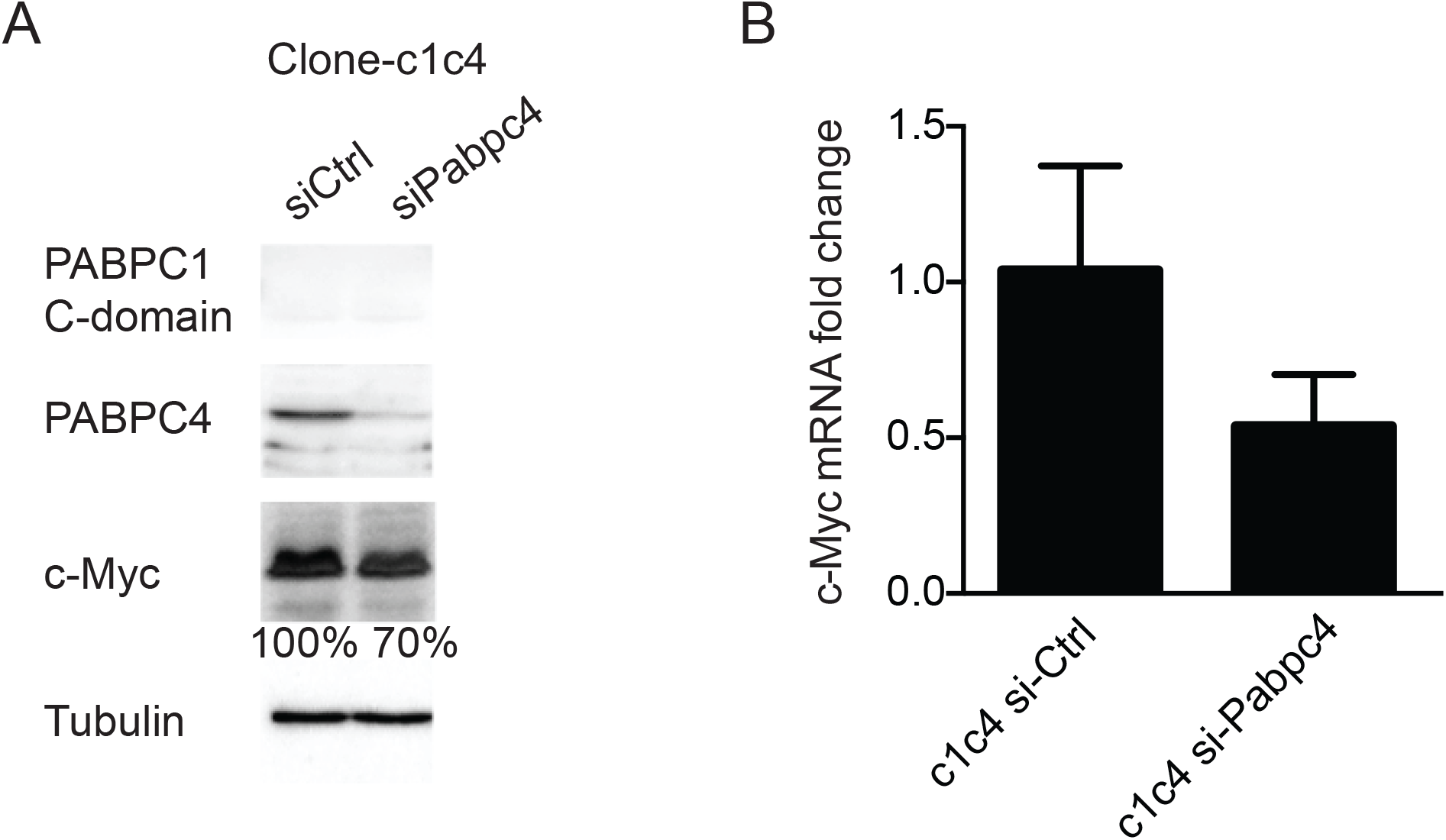
(A) Decrease of c-Myc protein level correlates with depletion of PABPC4. Representative blots of *Pabpc4* knocked-down clone-c1c4 cells. Corresponding antibodies were used to probe PABPC1, PABPC4, c-Myc, and tubulin levels. (B) Correlated c-Myc mRNA decrease after siRNA mediated knock-down of PABPC4.

### 3.9 Isoform usage of PABP and c-Myc levels

To test the effects of PABP isoform usage on c-Myc levels, we depleted PABPC1 or PABPC4 with siRNAs in HEK293 cells (Fig. 8A). Depletions of the two isoforms affect c-Myc mRNA levels in opposite directions. Decrease of PABPC4 lowered c-Myc mRNA, while decrease of PABPC1 raised c-Myc mRNA level. This confirms that the usage of the PABPC1 or PABPC4 isoform can influence the transcriptome through c-Myc.

**Figure 8.**
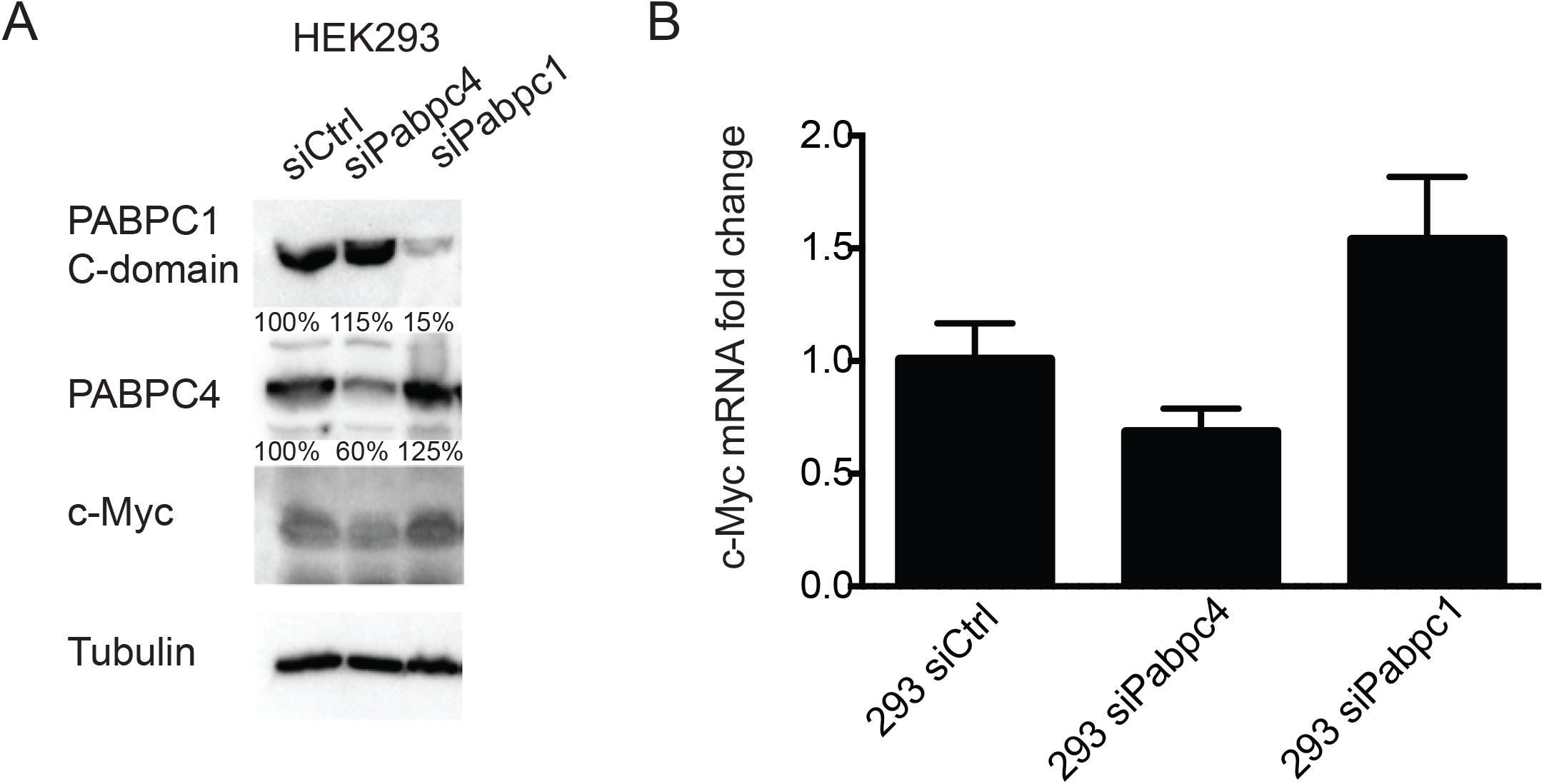
Differential effects of PABPC1 or PABPC4 depletion on c-Myc mRNA level in HEK293 cells. (A) HEK293 cells were transfected with control siRNA, si*Pabpc1*, or si*Pabpc4* for 48 hrs. PABPC1 and PABPC4 were probed to examine knock-down effects. (B) Changes of c-Myc mRNA levels in siRNA treated HEK293 cells. Depletion of PABPC4 reduces c-Myc mRNA level, while depletion of PABPC1 increases c-Myc mRNA. Both differences were significant at a p-value of 0.05 in a two-tail t-test for samples with identical variance.

## 4. Discussion

Here, we reported compensation of PABPC4 for the partial loss of PABPC1 in HEK293 cells. The cells are viable suggesting a functional overlap of the two isoforms. Analysis of the transcriptome profile revealed differential gene expression correlated with usage of PABPC4 and PABPC1. The study helps us understand isoform specific functions of PABP in development or different tissues.

Recent genomic studies support PABPC1 recognition of RNA sequences other than pure poly(A) in mouse (Kini, Silverman et al. 2016) and yeast (Baejen, Torkler et al. 2014). The overall structures of RRM domains in PABP isoforms are likely to be similar, due to high sequence similarities between domains. However, conserved differences are found in PABP isoforms across species. Some of the differences are located at interfaces critical for RNA recognition, especially in the RRM3-4 domains (Fig. S5). These differences do not alter the overall structure of thee RRM domains, but may contribute to specificity in RNA recognition. There is a growing realization that PABP isoforms are functionally different in vertebrate development (Gorgoni, Richardson et al. 2011) and other contexts. PABPC4 depletion impacts steady-state expression of a subset of mRNAs and affects erythroid differentiation (Kini, Kong et al. 2014). PABPC1L (ePABP) regulates translation and stability of maternal mRNAs (Vasudevan, Seli et al. 2006), and is substituted by PABPC1 after onset of zygotic transcription (Cosson, Couturier et al. 2002).

The recently developed *c*ross*l*inking *i*mmuno*p*recipitation coupled high-throughput *seq*uencing technique (CLIP-seq) provides a powerful tool to map association of RNA-binding proteins with RNA. Mapping the interactions of PABP isoforms with mRNA would greatly facilitate understanding of the functions of those isoforms. It is likely that PABP isoforms function through recognition of separate subsets of mRNAs, besides binding to a common mRNA pool. So far, such mapping data is only available for PABPC1 in higher eukaryotes (Kini, Silverman et al. 2016). The lack of CLIP data from other PABP isoforms makes biological inferences of differential gene expression profiles difficult. In this study, we identified c-Myc as a central player in remodeling the transcriptome of clone-c1c4 cells. C-Myc is a transcription factor that can shape the cellular transcriptome (Kress, Sabo et al. 2015). PABP isoforms may act through a subset of mRNAs to indirectly upregulate c-Myc transcription. In the Gene Set Enrichment Analysis, WNT pathway is enriched (Fig. 5) and can induce c-Myc transcription (Kress, Sabo et al. 2015). Mechanisms underlying correlation of PABPC4 and c-Myc remain to be revealed by further studies.

The significant increase (10-fold) of c-Myc mRNA is tolerated in clone-c1c4. Cell cycle analysis by propidium iodide DNA staining and flow cytometry showed similar distribution in HEK293 and clone-c1c4 cells in different phases of the cell cycle (data not shown). This may be because the increase in c-Myc protein levels were smaller, approximately three-fold. Elevation of genes like Axin1 (Supplemental Table 1) may enhance c-Myc protein turnover (Arnold, Zhang et al. 2009), which balances the sharp increase of c-Myc mRNA. Nonetheless, c-Myc targets are upregulated in clone-c1c4 extensively (Fig. 5B, Supplemental Tables 3 & 4 GSEA sets). Variant calling on the RNA-seq data reveals no insertion, deletion or mutation in c-Myc, or genes we know to affect c-Myc transcription. Blast search returns no other genomic sequence for the first 13 base pairs of the target sequence 2 (Fig. S1), which was used for generation of clone-c1c4. Modulation of PABPC4 or PABPC1 levels support a correlation of PABPC4 and c-Myc levels (Fig. 7&8). One interesting observation is that PABPC4 overexpression in HEK293 decreased endogenous PABPC1 by ∼50% (Fig. 3B), but did not affect the level of c-Myc mRNA (data not shown). The may reflect the dominant role PABPC1 compared to other PABP isoforms. Other isoforms can only function significantly when PABPC1 is at very low levels, as in clone-c1c4 (Fig. 7A) or PABPC1 depleted HEK293 (Fig. 8A). In summary, we created a human cell-line where the predominant PABPC1 is stably substituted by PABPC4. This opens a window to observe functional differences between the two isoforms. Validations and further investigations in different approaches will help us understand the mechanistic details of PABP regulation.

## Acknowledgements

We thank Dr. Guennadi Kozlov for critical comments. We are grateful for suggestions and discussion from colleagues during the project.

## Author contributions

J.X. designed and carried out the experiments. X.W. and Y.C. assisted J.X. with experiments. J.X. did the bioinformatics analysis and wrote the manuscript. K.G. revised the manuscript.

