## Supplemental figures for "Loss of PABPC1 is compensated by elevated PABPC4 and correlates with transcriptome changes"

PABPC1 genomic locus

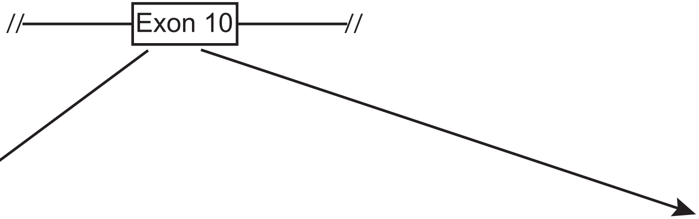


| | |
|---|---|
| PABPC1 genomic sequence | AGCATTCCAAAATATGCCCGGTGCTATCCGCCCAGCTGCTCCTAGACCACCA |
| Target 1 | GGTGCTATCCGCCCAGCTGC |
| Target 2 | AAATATGCCCGGTGCTATCC |

A

gBlocks sequence for target site 1:

TGTACAAAAAAGCAGGCTTTAAAGGAACCAATTCAGTCGACTGGATCCGGTACC
AAGGTCGGGCAGGAAGAGGGCCTATTTCCCATGATTCCTTCATATTTGCATATA
CGATACAAGGCTGTTAGAGAGATAATTAGAATTAATTTGACTGTAAACACAAAG
ATATTAGTACAAAATACGTGACGTAGAAAGTAATAATTTCTTGGGTAGTTTGCA
GTTTTAAAATTATGTTTTAAAATGGACTATCATATGCTTACCGTAACTTGAAAG
TATTTCGATTTCTTGGCTTTATATATCTTGTGGAAAGGACGAAACACC**GCAGCT**
**GGGCGGATAGCACC**GTTTTAGAGCTAGAAATAGCAAGTTAAAATAAGGCTAGTC
CGTTATCAACTTGAAAAAGTGGCACCGAGTCGGTGCTTTTTTTCTAGACCCAGC
TTTCTTGTACAAAGTTGGCATTA

gBlocks sequence for target site 2:

TGTACAAAAAAGCAGGCTTTAAAGGAACCAATTCAGTCGACTGGATCCGGTACC
AAGGTCGGGCAGGAAGAGGGCCTATTTCCCATGATTCCTTCATATTTGCATATA
CGATACAAGGCTGTTAGAGAGATAATTAGAATTAATTTGACTGTAAACACAAAG
ATATTAGTACAAAATACGTGACGTAGAAAGTAATAATTTCTTGGGTAGTTTGCA
GTTTTAAAATTATGTTTTAAAATGGACTATCATATGCTTACCGTAACTTGAAAG
TATTTCGATTTCTTGGCTTTATATATCTTGTGGAAAGGACGAAACACC**GGATAG**
**CACCGGGCATATTT**GTTTTAGAGCTAGAAATAGCAAGTTAAAATAAGGCTAGTC
CGTTATCAACTTGAAAAAGTGGCACCGAGTCGGTGCTTTTTTTCTAGACCCAGC
TTTCTTGTACAAAGTTGGCATTA

B

AAGAATTTTGGAGAAGACATGGATGATGAGCGCCTTAAGGATCTCTTTGGCAAGTTT
GGGCCTGCCTTAAGTGTGAAAGTAATGACTGATGAAAGTGGAAAATCCAAAGGATTT
GGATTTGTAAGCTTTGAAAGGCATGAAGATGCACAGAAAGCTGTGGATGAGATGAAC
GGAAAGGAGCTCAATGGAAAACAAATTTATGTTGGTCGAGCTCAGAAAAAGGTGGAA
CGGCAGACGGAACTTAAGCGCAAATTTGAACAGATGAAACAAGATAGGATCACCAGA
TACCAGGGTGTTAATCTTTATGTGAAAAATCTTGATGATGGTATTGATGATGAACGT
CTCCGGAAAGAGTTTTCTCCATTTGGTACAATCACTAGTGCAAAGGTTATGATGGAG
GGTGGTCGCAGCAAAGGGTTTGGTTTTGTATGTTTCTCCTCCCCAGAAGAAGCCACT
AAAGCAGTTACAGAAATGAACGGTAGAATTGTGGCCACAAAGCCATTGTATGTAGCT
TTAGCTCAGCGCAAAGAAGAGCGCCAGGCTCACCTCACTAACCAGTATATGCAGAGA
ATGGCAAGTGTACGAGCTGTTCCCAACCCTGTAATCAACCCCTACCAGCCAGCACCT
CCTTCAGGTTACTTCATGGCAGCTATCCCACAGACTCAGAACCGTGCTGCATACTAT
CCTCCTAGCCAAATTGCTCAACTAANACCAAGTCCTCGCTGGACTGCTCAGGGTGCC
AGACCTCATCC--------TAACACATCAACACAGACAATGGGTCCACGTCCTGCAG
CTGCAGCCGCTGCAGCTACTCCTGCTGTCCGCACCGTTCCACAGTATAAATATGCTG
CAGGAGTTCGCAATCCTCAGCNACATCTTAATGCACAGCCNCAAGTTACNATGCAAC
AGCCTGCTGTTCATGTACAAGGTCAGGAACCTTTGACTGCTTNCATGNTGNCNTCTG
CCCCTCCTCANAGCAAAAGCAAATGNNGNNGAACGGCT

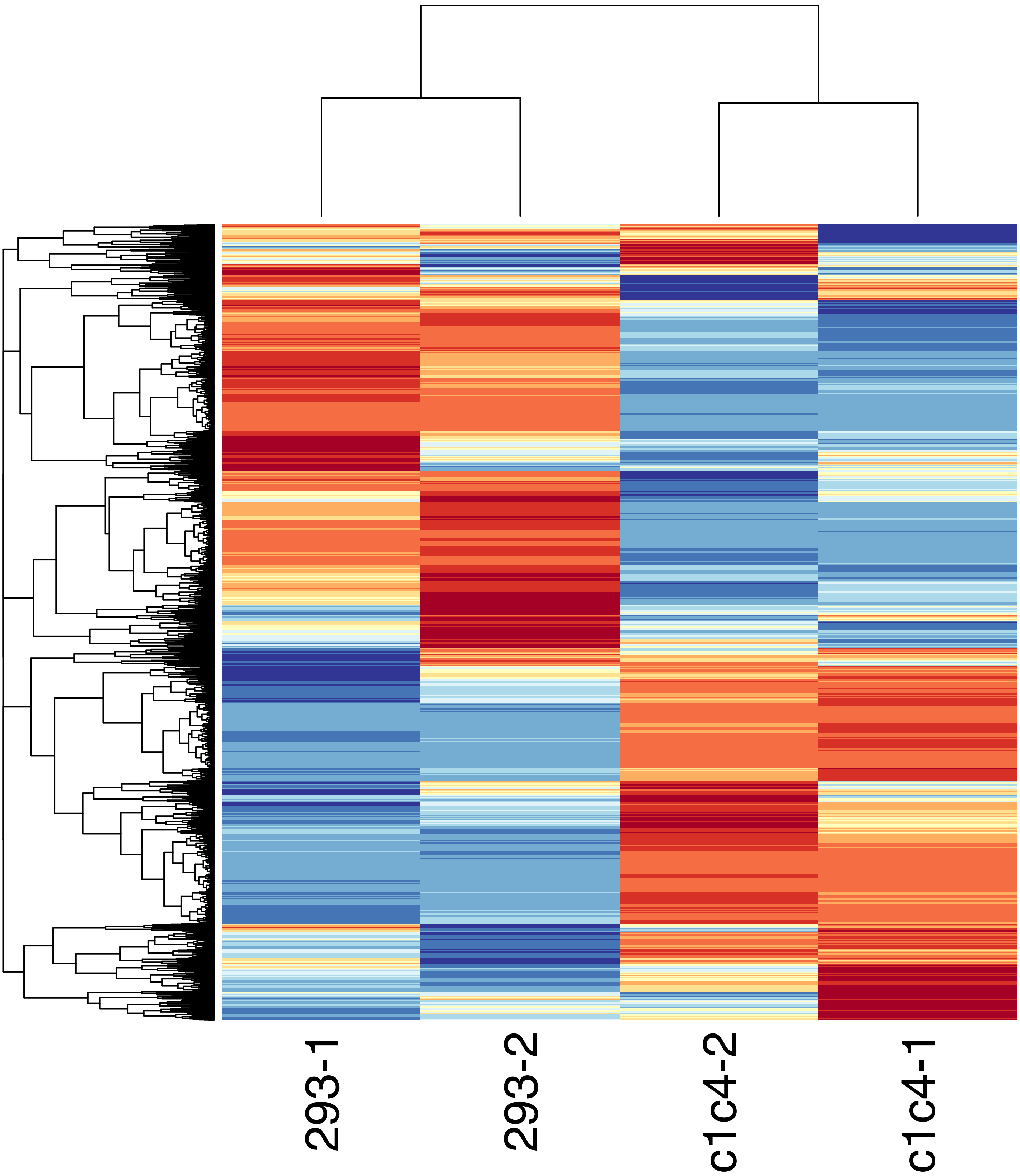

293-1
293-2
c1c4-2
c1c4-1

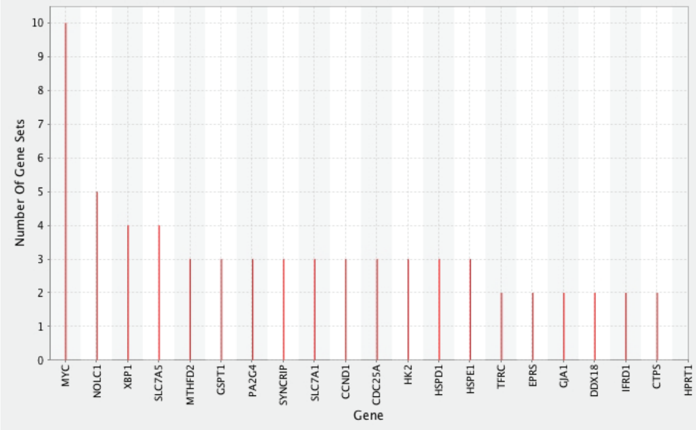
Number Of Gene Sets
0
1
2
3
4
5
6
7
8
9
10
MYC
NOLC1
XBP1
SLC7A5
MTHFD2
GSPT1
PA2G4
SYNCRIP
SLC7A1
CCND1
CDC25A
HK2
HSPD1
HSPE1
TFRC
EPRS
GJA1
DDX18
IFRD1
CTPS
HPRT1
Gene

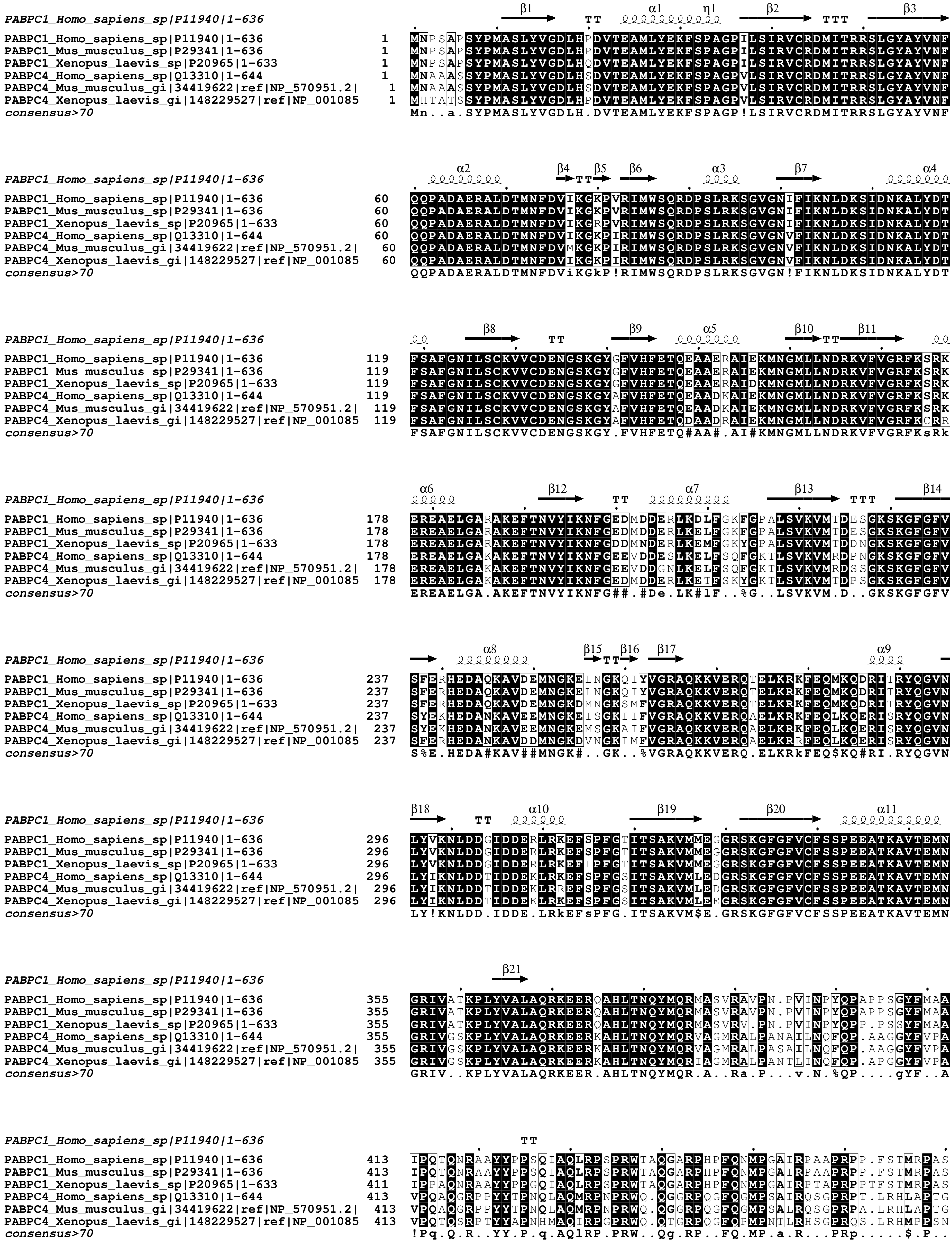
PABPC1_Homo_sapiens_sp|P11940|1-636
β1 TT α1 η1 β2 TTT β3
PABPC1_Homo_sapiens_sp|P11940|1-636 1 MNPSAPSYPMASLYVGDLHPDVTEAMLYEKFSPAGPILSIRVCRDMITRRSLGYAYVNF
PABPC1_Mus_musculus_sp|P29341|1-636 1 MNPSAPSYPMASLYVGDLHPDVTEAMLYEKFSPAGPILSIRVCRDMITRRSLGYAYVNF
PABPC1_Xenopus_laevis_sp|P20965|1-633 1 MNPSAPSYPMASLYVGDLHQDVTEAMLYEKFSPAGPILSIRVCRDMITRRSLGYAYVNF
PABPC4_Homo_sapiens_sp|Q13310|1-644 1 MNAAASSYPMASLYVGDLHSDVTEAMLYEKFSPAGPVLSIRVCRDMITRRSLGYAYVNF
PABPC4_Mus_musculus_gi|34419622|ref|NP_570951.2| 1 MNAAASSYPMASLYVGDLHSDVTEAMLYEKFSPAGPVLSIRVCRDMITRRSLGYAYVNF
PABPC4_Xenopus_laevis_gi|148229527|ref|NP_001085 1 MHTATSSYPMASLYVGDLHPDVTEAMLYEKFSPAGPVLSIRVCRDMITRRSLGYAYVNF
consensus>70 Mn..a.SYPMASLYVGDLH.DVTEAMLYEKFSPAGP!LSIRVCRDMITRRSLGYAYVNF
PABPC1_Homo_sapiens_sp|P11940|1-636
α2 β4 TT β5 β6 α3 β7 α4
PABPC1_Homo_sapiens_sp|P11940|1-636 60 QQPADAERALDTMNFDVIKGKPVRIMWSQRDPSLRKSGVGNIFIKNLDKSIDNKALYDT
PABPC1_Mus_musculus_sp|P29341|1-636 60 QQPADAERALDTMNFDVIKGKPVRIMWSQRDPSLRKSGVGNIFIKNLDKSIDNKALYDT
PABPC1_Xenopus_laevis_sp|P20965|1-633 60 QQPADAERALDTMNFDVIKGRPVRIMWSQRDPSLRKSGVGNIFIKNLDKSIDNKALYDT
PABPC4_Homo_sapiens_sp|Q13310|1-644 60 QQPADAERALDTMNFDVIKGKPIRIMWSQRDPSLRKSGVGNVFIKNLDKSIDNKALYDT
PABPC4_Mus_musculus_gi|34419622|ref|NP_570951.2| 60 QQPADAERALDTMNFDVMKGKPIRIMWSQRDPSLRKSGVGNVFIKNLDKSIDNKALYDT
PABPC4_Xenopus_laevis_gi|148229527|ref|NP_001085 60 QQPADAERALDTMNFDVIKGKPIRIMWSQRDPSLRKSGVGNVFIKNLDKSIDNKALYDT
consensus>70 QQPADAERALDTMNFDViKGkP!RIMWSQRDPSLRKSGVGN!FIKNLDKSIDNKALYDT
PABPC1_Homo_sapiens_sp|P11940|1-636
β8 TT β9 α5 β10 TT β11
PABPC1_Homo_sapiens_sp|P11940|1-636 119 FSAFGNILSCKVVCDENGSKGYGFVHFETQEAAERAIEKMNGMLLNDRKVFVGRFKSRK
PABPC1_Mus_musculus_sp|P29341|1-636 119 FSAFGNILSCKVVCDENGSKGYGFVHFETQEAAERAIEKMNGMLLNDRKVFVGRFKSRK
PABPC1_Xenopus_laevis_sp|P20965|1-633 119 FSAFGNILSCKVVCDENGSKGYGFVHFETQEAAERAIDKMNGMLLNDRKVFVGRFKSRK
PABPC4_Homo_sapiens_sp|Q13310|1-644 119 FSAFGNILSCKVVCDENGSKGYAFVHFETQEAADKAIEKMNGMLLNDRKVFVGRFKSRK
PABPC4_Mus_musculus_gi|34419622|ref|NP_570951.2| 119 FSAFGNILSCKVVCDENGSKGYAFVHFETQEAADKAIEKMNGMLLNDRKVFVGRFKSRK
PABPC4_Xenopus_laevis_gi|148229527|ref|NP_001085 119 FSAFGNILSCKVVCDENGSKGYAFVHFETQDAADKAIEKMNGMLLNDRKVFVGRFKCRR
consensus>70 FSAFGNILSCKVVCDENGSKGY.FVHFETQ#AA#.AI#KMNGMLLNDRKVFVGRFKsRk
PABPC1_Homo_sapiens_sp|P11940|1-636
α6 β12 TT α7 β13 TTT β14
PABPC1_Homo_sapiens_sp|P11940|1-636 178 EREAELGARAKEFTNVYIKNFGEDMDDERLKDLFGKFGPALSVKVMTDESGKSKGFGFV
PABPC1_Mus_musculus_sp|P29341|1-636 178 EREAELGARAKEFTNVYIKNFGEDMDDERLKELFGKFGPALSVKVMTDESGKSKGFGFV
PABPC1_Xenopus_laevis_sp|P20965|1-633 178 EREAELGARAKEFTNVYIKNFGDDMNDERLKEMFGKYGPALSVKVMTDDNGKSKGFGFV
PABPC4_Homo_sapiens_sp|Q13310|1-644 178 EREAELGAKAKEFTNVYIKNFGEEVDDESLKELFSQFGKTLSVKVMRDPNGKSKGFGFV
PABPC4_Mus_musculus_gi|34419622|ref|NP_570951.2| 178 EREAELGAKAKEFTNVYIKNFGEEVDDGNLKELFSQFGKTLSVKVMRDSSGKSKGFGFV
PABPC4_Xenopus_laevis_gi|148229527|ref|NP_001085 178 EREAELGAKAKEFTNVYIKNFGEDMDDERLKETFSKYGKTLSVKVMTDPSGKSKGFGFV
consensus>70 EREAELGA.AKEFTNVYIKNFG##.#De.LK#lF..%G..LSVKVM.D..GKSKGFGFV
PABPC1_Homo_sapiens_sp|P11940|1-636
α8 β15 TT β16 β17 α9
PABPC1_Homo_sapiens_sp|P11940|1-636 237 SFERHEDAQKAVDEMNGKELNGKQIYVGRAQKKVERQTELKRKFEQMKQDRITRYQGVN
PABPC1_Mus_musculus_sp|P29341|1-636 237 SFERHEDAQKAVDEMNGKELNGKQIYVGRAQKKVERQTELKRKFEQMKQDRITRYQGVN
PABPC1_Xenopus_laevis_sp|P20965|1-633 237 SFERHEDAQKAVDEMNGKDMNGKSMFVGRAQKKVERQTELKRKFEQMKQDRITRYQGVN
PABPC4_Homo_sapiens_sp|Q13310|1-644 237 SYEKHEDANKAVEEMNGKEISGKIIFVGRAQKKVERQAELKRKFEQLKQERISRYQGVN
PABPC4_Mus_musculus_gi|34419622|ref|NP_570951.2| 237 SYEKHEDANKAVEEMNGKEMSGKAIFVGRAQKKVERQAELKRKFEQLKQERISRYQGVN
PABPC4_Xenopus_laevis_gi|148229527|ref|NP_001085 237 SFERHEDANKAVDDMNGKDVNGKIMFVGRAQKKVERQAELKRRFEQLKQERISRYQGVN
consensus>70 S%E.HEDA#KAV##MNGK#..GK..%VGRAQKKVERQ.ELKRkFEQ$KQ#RI.RYQGVN
PABPC1_Homo_sapiens_sp|P11940|1-636
β18 TT α10 β19 β20 α11
PABPC1_Homo_sapiens_sp|P11940|1-636 296 LYVKNLDDGIDDERLRKEFSPFGTITSAKVMMEGGRSKGFGFVCFSSPEEATKAVTEMN
PABPC1_Mus_musculus_sp|P29341|1-636 296 LYVKNLDDGIDDERLRKEFSPFGTITSAKVMMEGGRSKGFGFVCFSSPEEATKAVTEMN
PABPC1_Xenopus_laevis_sp|P20965|1-633 296 LYVKNLDDGIDDERLRKEFLPFGTITSAKVMMEGGRSKGFGFVCFSSPEEATKAVTEMN
PABPC4_Homo_sapiens_sp|Q13310|1-644 296 LYIKNLDDTIDDEKLRKEFSPFGSITSAKVMLEDGRSKGFGFVCFSSPEEATKAVTEMN
PABPC4_Mus_musculus_gi|34419622|ref|NP_570951.2| 296 LYIKNLDDTIDDEKLRREFSPFGSITSAKVMLEDGRSKGFGFVCFSSPEEATKAVTEMN
PABPC4_Xenopus_laevis_gi|148229527|ref|NP_001085 296 LYIKNLDDTIDDEKLRKEFSPFGSITSAKVMLEEGRSKGFGFVCFSSPEEATKAVTEMN
consensus>70 LY!KNLDD.IDDE.LRkEFsPFG.ITSAKVM$E.GRSKGFGFVCFSSPEEATKAVTEMN
PABPC1_Homo_sapiens_sp|P11940|1-636
β21
PABPC1_Homo_sapiens_sp|P11940|1-636 355 GRIVATKPLYVALAQRKEERQAHLTNQYMQRMASVRAVPN.PVINPYQPAPPSGYFMAA
PABPC1_Mus_musculus_sp|P29341|1-636 355 GRIVATKPLYVALAQRKEERQAHLTNQYMQRMASVRAVPN.PVINPYQPAPPSGYFMAA
PABPC1_Xenopus_laevis_sp|P20965|1-633 355 GRIVATKPLYVALAQRKEERQAHLTNQYMQRMASVRV.PN.PVINPYQPP.PSSYFMAA
PABPC4_Homo_sapiens_sp|Q13310|1-644 355 GRIVGSKPLYVALAQRKEERKAHLTNQYMQRVAGMRALPANAILNQFQP.AAGGYFVPA
PABPC4_Mus_musculus_gi|34419622|ref|NP_570951.2| 355 GRIVGSKPLYVALAQRKEERKAHLTNQYMQRVAGMRALPASAILNQFQP.AAGGYFVPA
PABPC4_Xenopus_laevis_gi|148229527|ref|NP_001085 355 GRIVGSKPLYVALAQRKEERKAHLTNQYMQRIAGMRALPANTLINQFQP.APGGYFVPA
consensus>70 GRIV..KPLYVALAQRKEER.AHLTNQYMQR.A..Ra.P...v.N.%QP....gYF..A
PABPC1_Homo_sapiens_sp|P11940|1-636
TT
PABPC1_Homo_sapiens_sp|P11940|1-636 413 IPQTQNRAAYYPPSQIAQLRPSPRWTAQGARPHPFQNMPGAIRPAAPRPP.FSTMRPAS
PABPC1_Mus_musculus_sp|P29341|1-636 413 IPQTQNRAAYYPPSQIAQLRPSPRWTAQGARPHPFQNMPGAIRPAAPRPP.FSTMRPAS
PABPC1_Xenopus_laevis_sp|P20965|1-633 411 IPPAQNRAAYYPPGQIAQLRPSPRWTAQGARPHPFQNMPGAIRPTAPRPPTFSTMRPAS
PABPC4_Homo_sapiens_sp|Q13310|1-644 413 VPQAQGRPPYYTPNQLAQMRPNPRWQ.QGGRPQGFQGMPSAIRQSGPRPT.LRHLAPTG
PABPC4_Mus_musculus_gi|34419622|ref|NP_570951.2| 413 VPQAQGRPPYYTPNQLAQMRPNPRWQ.QGGRPQGFQGMPSALRQSGPRPA.LRHLAPTG
PABPC4_Xenopus_laevis_gi|148229527|ref|NP_001085 413 VPQTQSRPTYYAPNHMAQIRPGPRWQ.QTGRPQGFQPMPNTLRHSGPRQS.LRHMPPSN
consensus>70 !Pq.Q.R..YY.P.q.AQlRP.PRW..Qg.RP..FQ.MP.a.R...PRp.....$.P..

```
PABPC1_Homo_sapiens_sp|P11940|1-636

PABPC1_Homo_sapiens_sp|P11940|1-636                    471  ................SQVP.RV.M......S.....TQRVANTSTQTMGPRPAAAAAA
PABPC1_Mus_musculus_sp|P29341|1-636                    471  ................SQVP.RV.M......S.....TQRVANTSTQTMGPRPAAAAAA
PABPC1_Xenopus_laevis_sp|P20965|1-633                  470  ................NQVP.RV.M......S.....AQRVANTSTQTMGPRPTTAAAA
PABPC4_Homo_sapiens_sp|Q13310|1-644                    470  ................SECPDRLAMDFGGAGAAQQGLTDSCQSGGVPTAVQNLAPRAAV
PABPC4_Mus_musculus_gi|34419622|ref|NP_570951.2|       470  NAPASRGLPTTAQRVGSECPDRLAMDFGGAGAAQQGLTDSCQSGGVPTAVPNLAPRAAV
PABPC4_Xenopus_laevis_gi|148229527|ref|NP_001085       470  ................A..............QGTRGIPGVTQRVGVSSSTQTMGPRPPV
consensus>70                                                 .................q.p.r..m.............q........t........aa.

                                                                                                                α12    η2
PABPC1_Homo_sapiens_sp|P11940|1-636

PABPC1_Homo_sapiens_sp|P11940|1-636                    501  ATPAVRTVPQYKYAAGVRNPQQHLNAQPQVTMQQPAVHVQGQEPLTASMLASAPPQEQK
PABPC1_Mus_musculus_sp|P29341|1-636                    501  ATPAVRTVPQYKYAAGVRNPQQHLNAQPQVTMQQPAVHVQGQEPLTASMLASAPPQEQK
PABPC1_Xenopus_laevis_sp|P20965|1-633                  500  AASAVRAVPQYKYAAGVRNQ.QHLNTQPQVAMQQPAVHVQGQEPLTASMLAAAPPQEQK
PABPC4_Homo_sapiens_sp|Q13310|1-644                    513  AAAAPRAVAPYKYAASSVRSPHP..AIQPL.QAPQPAVHVQGQEPLTASMLAAAPPQEQK
PABPC4_Mus_musculus_gi|34419622|ref|NP_570951.2|       529  AAAAPRAVAPYKYAASSVRSPHP..AIQPL.QAPQPAVHVQGQEPLTASMLAAAPPQEQK
PABPC4_Xenopus_laevis_gi|148229527|ref|NP_001085       499  SAPPPRAVPPYKYT...RCPLP..VVQPL.QAPQPAVHVQGQEPLTASMLASAPPQEQK
consensus>70                                                 a..a.R.V..YKYa..vR.p......QP.....QPAVHVQGQEPLTASMLA.APPQEQK

                                                               α13        α14         η3       α15                α16
PABPC1_Homo_sapiens_sp|P11940|1-636

PABPC1_Homo_sapiens_sp|P11940|1-636                    560  QMLGERLFPLIQAMHPTLAGKITGMLLEIDNSELLHMLESPESLRSKVDEAVAVLQAHQ
PABPC1_Mus_musculus_sp|P29341|1-636                    560  QMLGERLFPLIQAMHPSLAGKITGMLLEIDNSELLHMLESPESLRSKVDEAVAVLQAHQ
PABPC1_Xenopus_laevis_sp|P20965|1-633                  558  QMLGERLFPLIQAMHPTLAGKITGMLLEIDNSELLHMLESPESLRSKVDEAVAVLQAHQ
PABPC4_Homo_sapiens_sp|Q13310|1-644                    569  QMLGERLFPLIQTMHSNLAGKITGMLLEIDNSELLHMLESPESLRSKVDEAVAVLQAHH
PABPC4_Mus_musculus_gi|34419622|ref|NP_570951.2|       585  QMLGERLFPLIQTMHSNLAGKITGMLLEIDNSELLHMLESPESLRSKVDEAVAVLQAHH
PABPC4_Xenopus_laevis_gi|148229527|ref|NP_001085       552  QMLGERLFPLIQAMHPSLAGKITGMLLEIDNSELLHMLESPESLRSKVEEAVAVLQAHQ
consensus>70                                                 QMLGERLFPLIQ.MH..LAGKITGMLLEIDNSELLHMLESPESLRSKV#EAVAVLQAH.

PABPC1_Homo_sapiens_sp|P11940|1-636

PABPC1_Homo_sapiens_sp|P11940|1-636                    619  AKEAAQKAVNSATGVPTV
PABPC1_Mus_musculus_sp|P29341|1-636                    619  AKEAAQKAVNSATGVPTV
PABPC1_Xenopus_laevis_sp|P20965|1-633                  617  AKEAAQKVVN.ATGVPTA
PABPC4_Homo_sapiens_sp|Q13310|1-644                    628  AKKEAAQKVGAVAA.ATS
PABPC4_Mus_musculus_gi|34419622|ref|NP_570951.2|       644  AKKEAAQKVGTVAA.ATS
PABPC4_Xenopus_laevis_gi|148229527|ref|NP_001085       611  AKKDAAQKVGIVT..ATS
consensus>70                                                 AK..A...V.......T.
```
